# Dynamic energy budget model for a bumble bee colony: Predicting the spatial distribution and dynamics of colonies across multiple seasons

**DOI:** 10.1101/2024.03.05.583508

**Authors:** Pau Capera-Aragones, Joany Mariño, Amy Hurford, Rebecca C. Tyson, Eric Foxall

## Abstract

Bumble bees are important pollinators of many crops around the world. In recent decades, agricultural intensification has resulted in significant declines in bumble bee populations and the pollination services they provide. Empirical studies have shown that this trend can be reversed, however, by enhancing the agricultural landscape with natural habitat, such as adding wildflower patches adjacent to crops. Despite the empirical evidence, the mechanisms behind these positive effects are not fully understood, and the specific characteristics of the enhanced natural habitat that would maximize benefits are unclear at this time. Theoretical studies, in the form of mathematical models, have proven useful in elucidating the underlying mechanisms and determining the optimal natural habitat configurations. Existing models, however, generally focus only on particular aspects of bumble bee behaviour; some models are accurate at describing population dynamics, while others are accurate at describing their spatial distribution. In this work, we build a unique model coupling population dynamics, using a whole-colony Dynamic Energy Budget (DEB) approach, to a spatial distribution model based on the maximum energy principle. This coupling gives valuable new insights into the effects of spatial arrangements on population dynamics, and vice-versa. With our model, we answer questions such as when, how much, or what type of wildflower patches should be planted to maximize crop pollination services and minimize bee decline. We find that planting wildflowers that bloom before and after crop bloom is crucial to achieve high pollination services and preserving wild pollinator populations. We also find that small quantities of natural habitat are needed when the crop is nutritionally rich, but higher quantities are most beneficial when the crop is nutritionally deficient.

## 1 Introduction

Wild pollinators, such as bumble bees, provide important pollination services to crops around the world (Porto et al., 2020; Winfree et al., 2009). It is therefore concerning that wild bees, and the pollination services they provide, have declined in several regions (Biesmeijer et al., 2006; Potts et al., 2005; Vanbergen, Initiative, et al., 2013). Agricultural intensification, characterized by the high use of pesticides and fertilizers, increases in farm size, low proportion of natural habitat in the landscape, and simplified crop rotation (Goulson et al., 2008; Mola et al., 2021; Stoate et al., 2001), are considered to be the major reasons for pollinator habitat loss and the widespread decline of farmland biodiversity (Robinson and Sutherland, 2002).

Several empirical studies have shown that the addition of natural habitat adjacent to pollinator-dependent crops is an effective strategy to prevent population and diversity declines in wild pollinators (Blaauw and Isaacs, 2014; Buhk et al., 2018; Carvalheiro et al., 2011). A key example is the study by Blaauw and Isaacs (2014), which shows that wildflower strips planted adjacent to blueberry crops can increase crop pollination services and, at the same time, increase pollinator population size and diversity. The benefits observed in Blaauw and Isaacs (2014) can already be seen in the first year after the wildflowers are planted, but increase significantly over the course of the 4-year study.

Despite the empirical evidence in favor of using wildflower enhancements to increase crop pollination, the costs of these management strategies together with the lack of clarity on the optimal location, quantity, flower type, or flowering time of the wildflower enhancements diminishes the practical effectiveness of these strategies (Albrecht et al., 2020). Recently, useful insights have been provided by several modelling studies (Capera-Aragones, Foxall, et al., 2021, 2022; Carturan et al., 2023; Häussler et al., 2017; Lonsdorf et al., 2009; MacQueen et al., 2022; Twiston-Davies et al., 2021). However, none of these models are able to fully describe the population dynamics, the spatial dynamics, the balance between nutrition and energy needs of bees, and their interplay, all of which are important in determining the effect of wildflower patches on pollination services.

In this work, we develop a novel mathematical model that accounts for both the population dynamics and the spatial distribution of wild bees, and we use the model to investigate the wildflower patch characteristics that prevent wild bee decline and provide the most benefit to crop pollination services. In our model, spatial dynamics is incorporated through application of the Maximum Entropy Principle as recently developed by Capera-Aragones, Tyson, et al. (2023). Population dynamics is incorporated using a Dynamic Energy Budget (DEB) framework, which is applied to the whole colony rather than to individuals as is normally the case. DEB models have been widely used in recent years to study a wide range of organisms (Humphries et al., 2004; Mariño et al., 2019; Pouvreau et al., 2006; Van Haren and Kooijman, 1993), including bumble bees (Kenna et al., 2019), but in all cases, the model is based on the energy budgets of the individuals, not on that of a whole colony. Applying the approach to the entire colony allows us to track the evolution of colony properties with minimum complexity, and at the same time increases the predictive capabilities of the model, allowing us to obtain colony-level insights about energy budget and management. Our model is the first to apply DEB at a colony level and to couple it to a spatial model.

With our approach, we arrive at a model that accounts for important ecological properties of the flower-pollinator system and yet has remarkable mathematical simplicity compared to other recent models (M. Becher et al., 2016; Matthias A Becher et al., 2018; Carturan et al., 2023). Our results show the importance of wildflower patches near to nutritionally deficient crops, such as blueberry and cranberry (Delaplane et al., 2000; Kremen et al., 2002; Richards, 2001), in line with earlier predictions Capera-Aragones, Foxall, et al. (2022). In addition, our results show the importance of considering the bloom time of the wildflower enhancements in relation to the bloom time of the crop. In particular, the benefits of added wildflowers are maximized when wildflowers bloom before the crop (to increase the pollinator population size before crop bloom), and after the crop blooms (to maintain a large number of hibernating queens over the seasons).

## 2 Model and Methods

Dynamic Energy Budget (DEB) models provide a quantitative framework to dynamically describe the energy and mass budgets of living organisms at the individual level, based on assumptions about energy uptake, storage, and utilization of various substances (Jusup et al., 2017; Meer, 2006; Sousa, Domingos, and Kooijman, 2008). DEB models use thermodynamic principles such as conservation of mass, energy and time, and relationships between the surface and the volume of the individuals to link different levels of biological organization, i.e. from cells to individuals to populations (Kooijman, 2010; Sousa, Domingos, Poggiale, et al., 2010).

The theory has been successfully applied to over 1000 different species (Marques et al., 2018) with applications in general ecology, conservation, aquaculture, and ecotoxicology (Lavaud et al., 2021). In our work, we develop a DEB-based model applied to the case of wild bees. In contrast to traditional DEB models, we consider the individual to be the whole colony of wild bees, and apply the DEB approach to that “super-organism”. This framing means that the energy fluxes in our model are those of the colony and not the individual bees. The energy uptake, for example, is not proportional to the nectar collected by an individual bee. Instead, it is proportional to the nectar collected by all bees, which is related to their foraging spatial distribution rather than the surface of an individual bee. Applying the DEB framework to the whole colony increases the predictive capabilities of the model at the colony level, and allows us to obtain insights into the energy budget and management of the whole colony.

The spatial distribution of foragers, i.e., where and when individual bees are able to find resources, has a strong effect on the energy fluxes of the colony. We therefore couple our DEB model to a novel spatially explicit model recently developed by Capera-Aragones, Tyson, et al. (2023). The spatial model can be understood as an extension of the Ideal Free Distribution (Fretwell, 1969) in which travelling costs, foraging efficiency, or resource depletion are not ignored. Using this approach to compute the spatial distribution of foragers keeps our model simple, while at the same time allowing us to gain realistic insights into the interaction between foragers and the landscape.

Fig. 1 is a graphical representation of how our approach tracks the energy fluxes of the colony to describe the evolution of four state variables: Nectar reserve (N), Pollen reserve (P), Structural energy (V), and Colony number (R). In Section 2.1 we describe the mathematical expressions for each of the energy fluxes involved in all vital functions of the colony. In Section 2.2 we describe the meaning and mathematical expressions for the evolution of each state variable. In Section 2.3 we describe the application of the Maximum Entropy Principle to compute the foraging spatial distributions. Finally, in section 2.4 we summarize and simplify the model, and we give the parameter values used for the numerical simulations.

**Figure 1:**
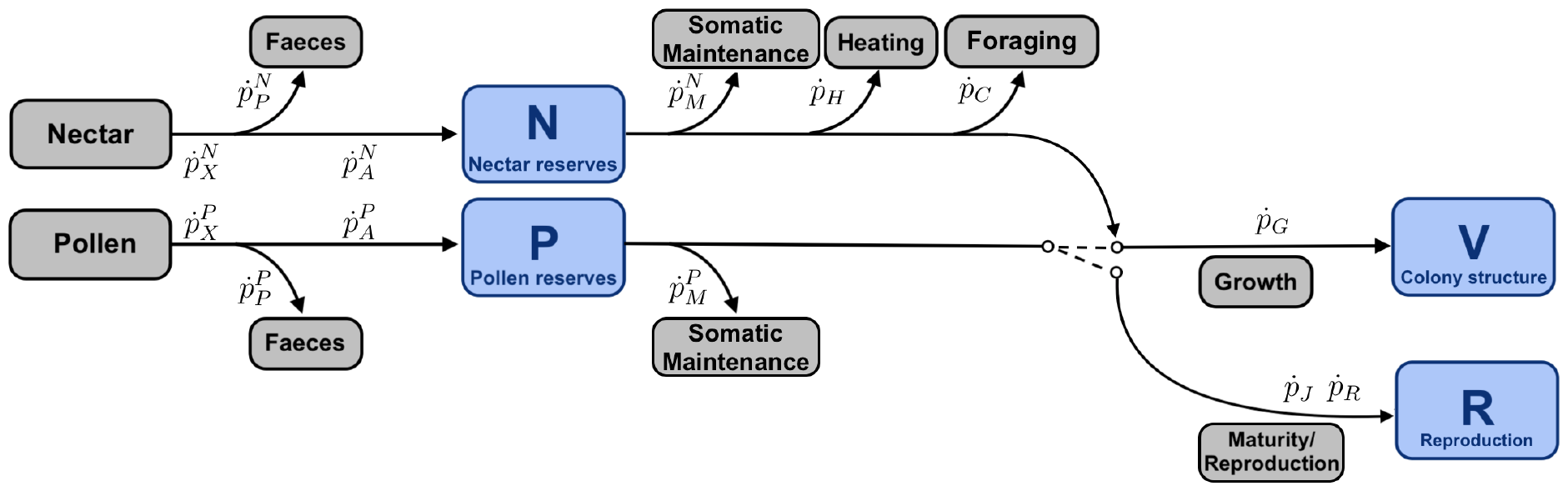
Diagram of the DEB model for a wild bumble bee colony. The switch represents a metabolic threshold at the end of the season. The four state variables are shown in blue.

### 2.1 Energy Fluxes

A basic step in the construction of a DEB model is the definition of the energy fluxes. The realism and complexity of the resulting model depends significantly on these definitions. The energy fluxes for our model are illustrated in Fig. 1, and defined below.

#### Ingestion (nectar 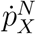and pollen 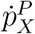)

We define

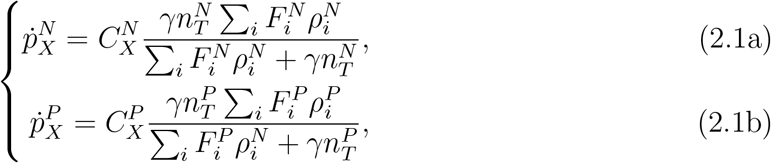

where 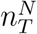 and 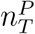 are the total number of nectar and pollen foragers respectively, 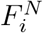 and 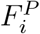 are the amount of pollen and nectar resources in patch *i* of the landscape, and 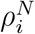 and 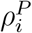 are the density of nectar and pollen foragers in the patch *i*. Ingestion is proportional to the consumption by the entire forager population (*γn*_*T*_) if the resource in the landscape is not limiting 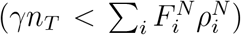, and it is proportional to the resource in the landscape 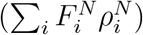 if the resource in the landscape is limiting 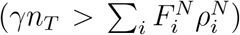. Increasing the number of foragers does not increase ingestion by the colony if there is no resource available in the landscape, and increasing the resource in the landscape does not increase ingestion if there are no additional foragers to collect it.

#### Faeces (nectar 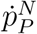 and pollen 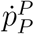 )

We define

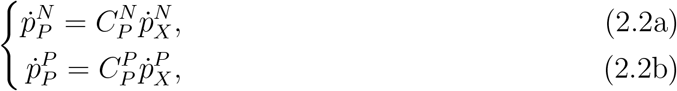

where 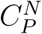 and 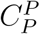 are constants of proportionality that dictate which part of the ingested food will not be incorporated into the reserves of the colony.

#### Assimilation (nectar 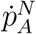 and pollen 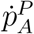 )

We define

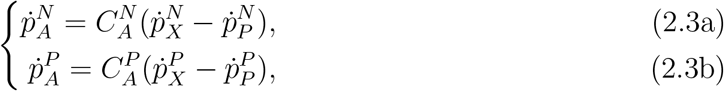

where 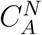 and 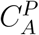 are proportionality constants that dictate what portion of the incorporated food will actually be digested.

#### Somatic maintenance (nectar 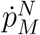 and pollen 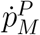 )

We define somatic maintenance as the process of expending energy to avoid or repair damage, to maintain concentration gradients of metabolites across membranes, and many other process that help preserving the integrity of the organism. Mathematically, we define

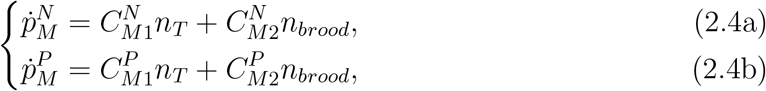

with *n*_*T*_ being the total number of foragers and *n*_*brood*_ the total brood population. 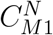 and 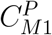 are constants of proportionality which determine the somatic maintenance associated to foragers, and 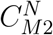 and 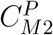 the ones associated to the somatic maintenance of the brood.

#### Heating 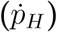

We define

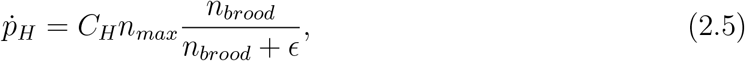

where *C*_*H*_ is a proportionality constant, *n*_*max*_ is the maximum size of the colony ever had during the course of the season (foragers and brood), and the fraction 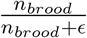is a proportional factor that deactivates the heating process (i.e. eliminates heating costs) in the absence of brood.

#### Foraging 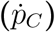

We define

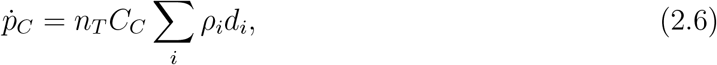

where *C*_*C*_ is a proportionality constant, *d*_*i*_ is the distance of the resource patch *i* relative to the nest, and 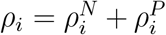is the probability distribution of nectar and pollen foragers. The sum in the previous equation depends on the forager spatial distribution in such a way that the foraging energy flux increases with forager distance from the nest.

#### Growth 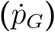

We define

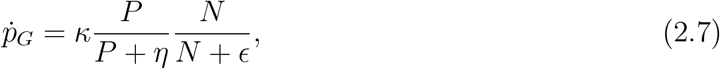

where *ϵ* is a small value that renders the growth independent of *N*, except in the case where *N* is very small and growth becomes limited by *N*. Similarly, colony growth is proportional to *P* when *P* is small, but saturates as *P* increases at a rate that depends on the rate *η* at which the queen bee lays eggs. *κ* is a proportionality constant that accounts for the amount of energy necessary to increase the structural volume.

#### Maturity maintenance 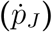

We define

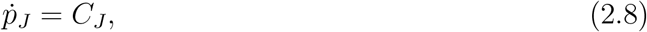

where *C*_*J*_ is simply a constant. This equation predicts a linear increase in maturity.

#### Reproduction/Maturation 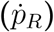

We define

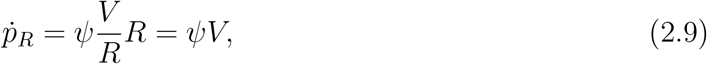

where *Ψ* is a proportionality constant, *V* is the structural energy of the colony (related to the number of foragers and brood) and *R* is the reproductive outcome (related to the number of colonies present in the landscape). Note that the colony reproduction is proportional to the per-colony structural volume (*V/R*) multiplied by the number of colonies (*R*), which simplifies to *ΨV* .

### 2.2 State Variables

As shown in Fig. 1, our model uses four state variables to describe the state of the dynamical system.

#### Nectar reserve (*N* )

This variable refers to the nectar stored in the nest. Nectar (which has a high concentration of sugar) is the main source of energy for the colony. Note that this state variable is separate from the nectar in the landscape 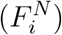.

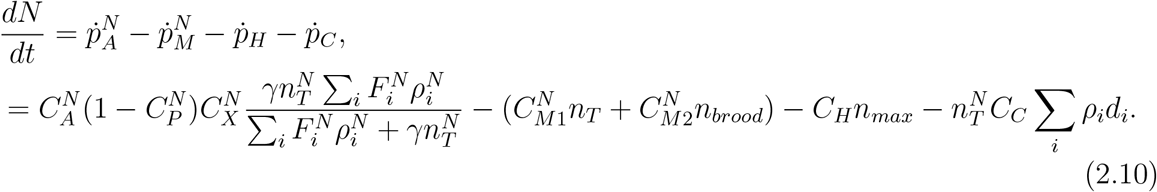

#### Pollen reserve (*P* )

This variable refers to the pollen stored in the nest. The pollen (which is largely composed of proteins) is the main source of amino acids used for body development. As with the nectar (*N*), this state variable is separate from the pollen in the landscape 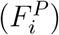.

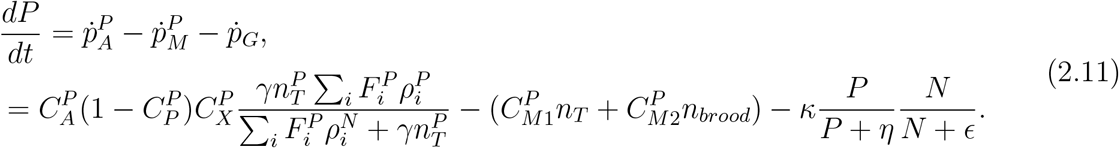

#### Structural energy (*V* )

This variable refers to the energy stored in the bodies of all foraging bees and brood.

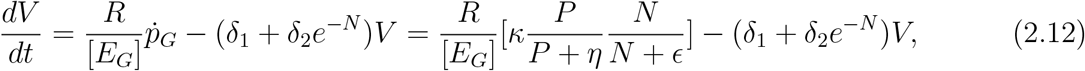

where [*E*_*G*_] is a measure of the rate of transformation of pollen into structural energy. This parameter is related to the rate at which queens lay eggs. If we want to consider multiple colonies, we need to multiply [*E*_*G*_] by the inverse of the number of colonies (*R*) (i.e., the number of queens producing eggs). *δ*_1_ is the “natural” death rate and 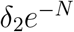 is an addition to the death rate that is negligible unless colony nectar stores become small.

#### Colony number (*R*)

This variable refers to the number of independent colonies (i.e., the number of queens) in the landscape, and so *R*∈ N. We do not consider the spatial distribution of the colonies in this version of the model, so we assume that the nest sites are concentrated in a single location. This is an important simplification of our model. By contrast to *N, P*, and *V*, the value of *R* only updates at the beginning of the season, and is described using a difference equation

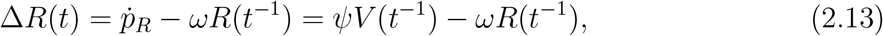

where *ΨV* is colony reproduction and *ωR*_*t−*1_ is colony death rate during winter.

### 2.3 Forager distribution (*n*)

To solve eqs. (2.10)-(2.13) we first need to find the forager spatial probability distribution, 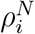 and 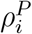, of nectar and pollen foragers, respectively, in a given landscape 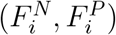, and for a given bee population *n*_*T*_. Finding spatial distribution of populations is a common problem in spatial ecology and can be done in many different ways. In our case, we will compute the spatial distribution using the Maximum Entropy principle as in Capera-Aragones, Tyson, et al. (2023). We obtain

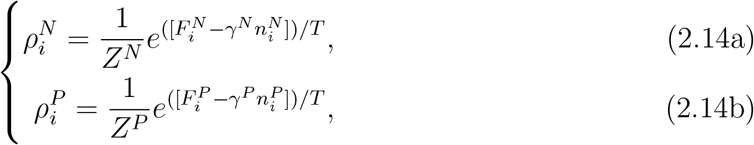

with *T* → 0 being optimal and *T* → ∞ corresponding to randomly distributed foragers. where *T* refers to the inverse of the optimality of the foraging strategy of individuals, *γ*^*N/P*^ are the pollen and nectar consumption rates and 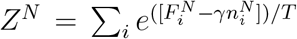 and 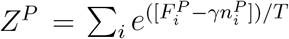 are normalization factors. The total number of nectar and pollen foragers (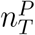and 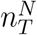) is given by

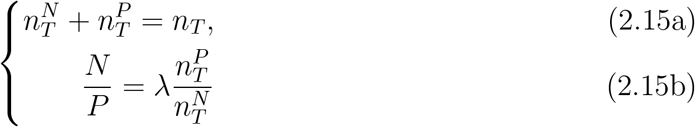

If we combine eq. (2.15) we obtain

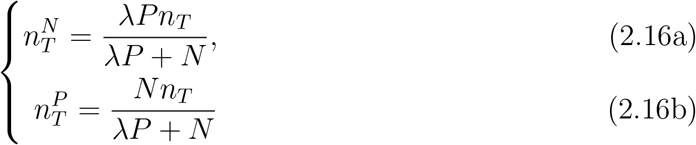

To make the model more realistic and numerically stable, Eqs. (2.16) need a correction factor. This factor is designed so each 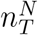 and 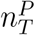 must represent at most 99% of the population and at least 1%. That is, there must be at least a few bees in each group of foragers. Hence, we have

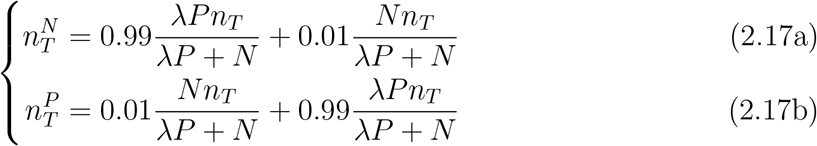

Using 2.14 and 2.17 we obtain the foraging spatial distributions that are needed to compute the temporal dynamics of the state variables:

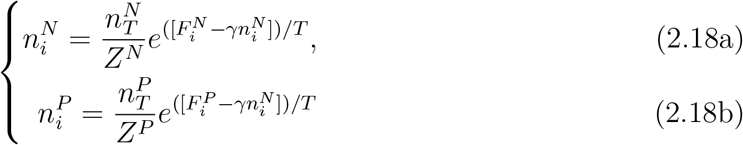

Our full model is defined by eqs. (2.10)-(2.13) with Eq.(2.19).

### 2.4 Multi-seasonality, simplifications and model summary

In this section we (1) extend the model to multi-seasonality, (2) make some simplifications, (3) summarize the model, and (4) give the parameter values used to perform the simulations.

Bumble bees have two major distinct behaviours throughout the year: (1) Worker bees forage around the landscape for the growth and survival of the colony and (2) Worker bees die and queen bees hibernate. In order to account for these two distinct behaviours, we divide the model into two phases:

#### Reproductive Impulse (for t ∈ℕ)

In spring, when the foraging season is about to start, the colony number is determined based on the success of the previous season, pollen and nectar reserves are set to zero, and the structural energy is set to a small value which accounts only for the presence of queen bees. The model at the beginning of the season is

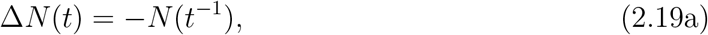

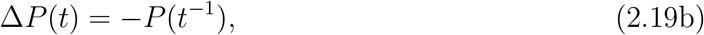

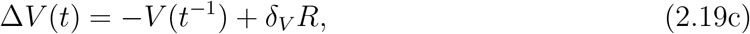

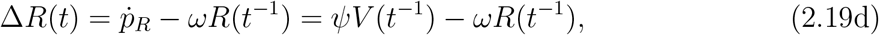

where *δ*_*V*_ *R* is the structural volume corresponding to all queen bees.

#### Colony growth (for t ∈ ℝ ⊃ ℕ)

Once the first set of working bees have matured and are foraging, colony growth proceeds according to eqs. (2.10)-(2.13). Assuming *n*_*T*_ = *αV* and *n*_*brood*_ = *βV*, defining 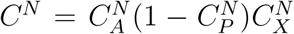 and 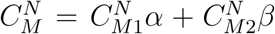 (similarly *C*^*P*^ and 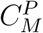), using eq. (2.16), the model throughout the season becomes

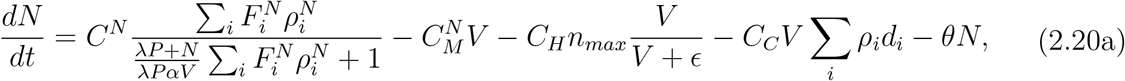

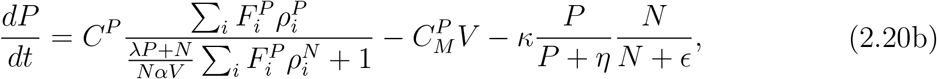

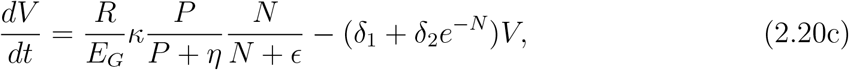

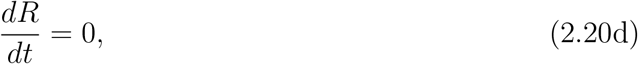

with the distributions of foragers given by

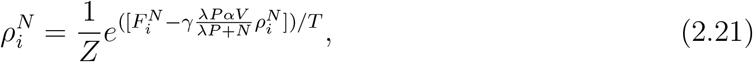

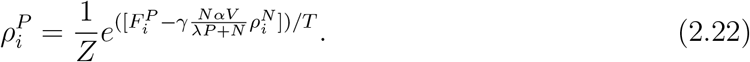

The nectar and pollen in the landscape are also dynamic and given by

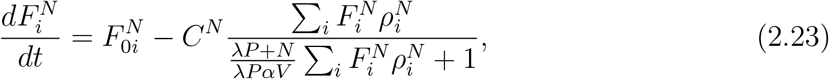

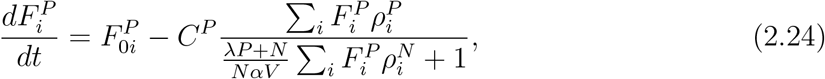

where the first term represents the production of nectar or pollen (which is related to the number of flowers in the region and is heterogeneous and dynamic in general), and the second term represents the ingestion of nectar or pollen by foragers.

A summary of the variables used in the model is given in Table.1. All parameter values are specified in Table.2. Parameter values are determined by requiring the model to correctly reproduce some expected qualitative behaviours. For example, *v* = 1 is the value that saturates the growth of the colony at the value of 50 individual bees, which is the typical average size observed for bumble bees in nature. In this work, we do not aim to reproduce quantitatively any particular experiment or field study; the focus of our analysis is to demonstrate the versatility of our approach, and the interesting qualitative behaviours that can be observed.

**Table 1:** Summary of variables and their meaning.

| Symbol | Meaning |
| --- | --- |
| $N$ | Nectar storage by all colonies |
| $P$ | Pollen storage by all colonies |
| $V$ | Structural volume (number of bees and brood) of all colonies |
| $R$ | Number of colonies |
| $\rho_i^N$ | Probability density of nectar foragers in patch $i$ |
| $\rho_i^P$ | Probability density of pollen foragers in patch $i$ |
| $F_i^N$ | Nectar in patch $i$ of the landscape |
| $F_i^P$ | Pollen in patch $i$ of the landscape |

**Table 2:**
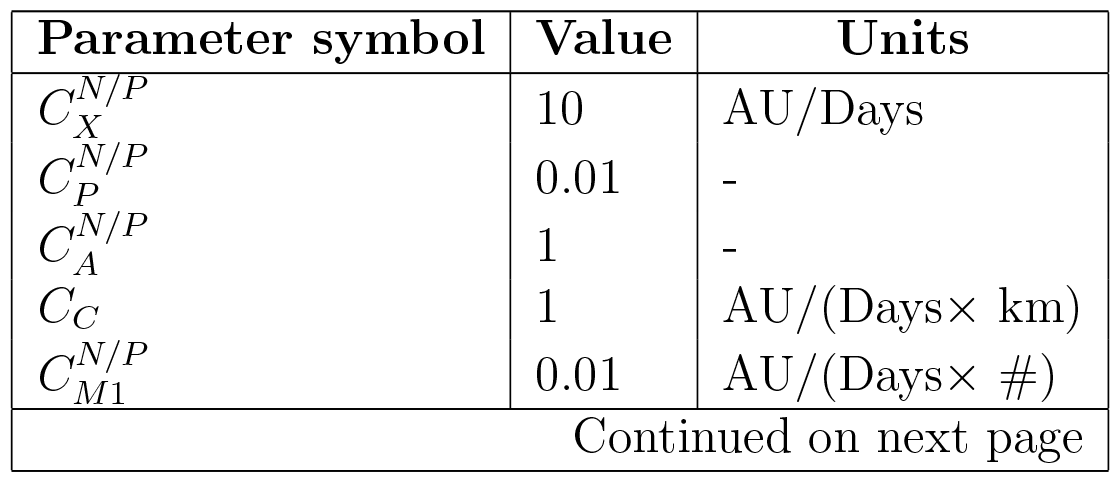
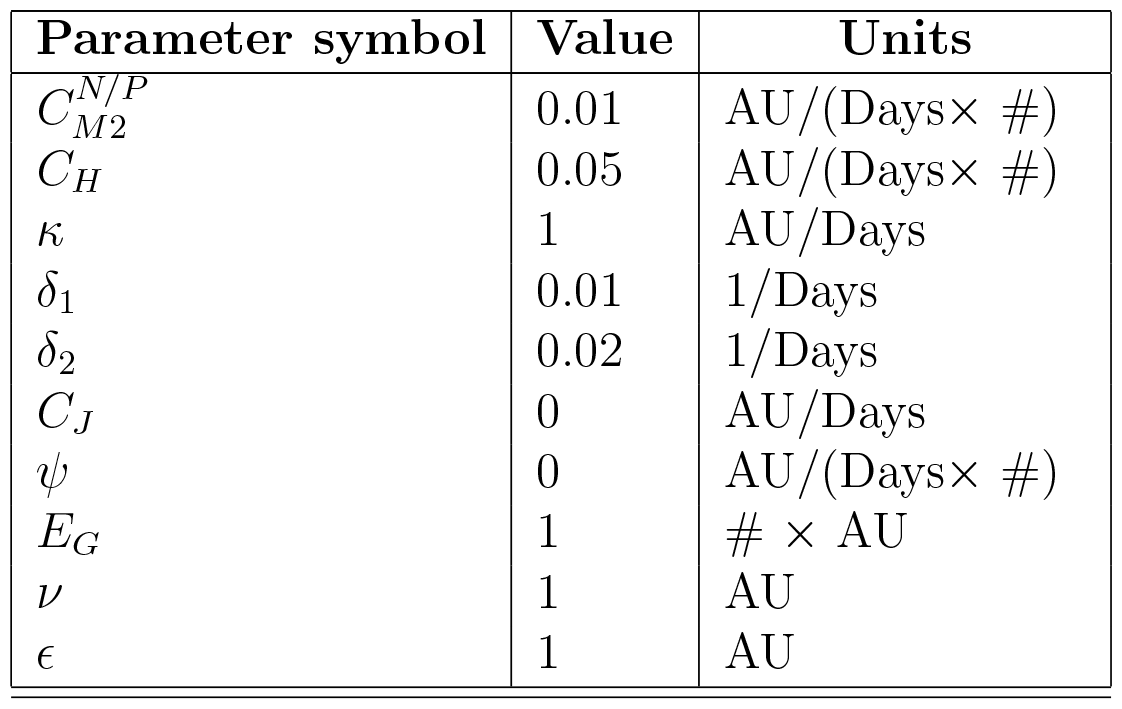
Summary of parameter values used for the simulations. ‘AU’ refers to ‘Arbitrary Units’ and ‘#’ refers to ‘number of bees’. The parameters of the models are set such that *time, distances*, and *bee populations* values are qualitatively correct, but *energy* and *resource quantity* values are arbitrary.

## 3 Results

We divide this section into two parts. In the first part, we focus on a single season and a single colony scenario, i.e, we do not consider the reproductive impulse (eqs. (2.19)) nor the state variable *R*, and we check the ability of the model to reproduce and predict behaviours that have been observed empirically or predicted by other existing models. In the second part, we study the ability of the model to reproduce and predict the behaviours observed in multiple seasons and multiple colony scenarios. In the second part, we also predict the location, quantity, type, and bloom time of the wildflower enhancements that will most benefit crop pollination services.

### 3.1 Single season and single colony

#### 3.1.1 Colony growth

We first investigate the effect of colony growth in a landscape with limited or unlimited resources. We simulated scenarios in which (1) pollen in the landscape is limited, (2) nectar in the landscape is limited, and (3) neither is limited. We then present the predicted population dynamics in each scenario. Fig. 2 shows that the growth of the colony can saturate in three different ways:

**Figure 2:**
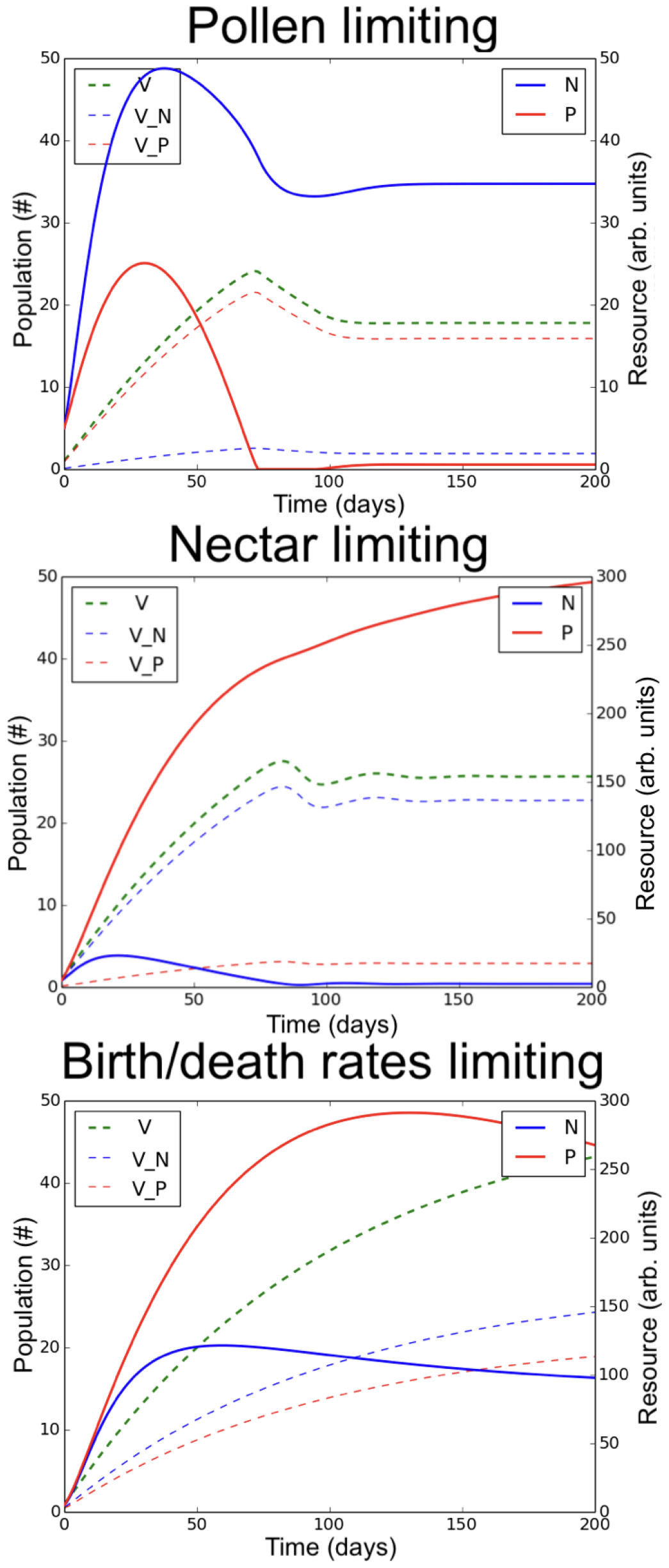
Structural energy of the colony (*V*) (stored in nectar foragers (*V*_*N*_) and pollen foragers (*V*_*P*_)), as well as nectar storage (*N*) and pollen storage (*P*) as functions of time, in a landscape with a uniform distribution of resource. In the first plot, pollen in the landscape is limiting, in the centre plot nectar is limiting, and in the bottom plot neither nectar nor pollen are limiting and only the birth and death rates limit population growth.

1. Limiting pollen (*P* ≈ *η*): The brood cannot be fed properly and the colony stops growing.
2. Limiting nectar (*N*≈ *ϵ*): Although brood requires mostly pollen for growth, a minimal quantity of nectar is also necessary for brood development. In addition, and most importantly, the absence of nectar results in increasing mortality of adult bees, since nectar is their main resource.
3. Limiting birth/mortality rate: When both pollen and nectar are abundant, the growth of the colony is limited by the rate at which the queen lay eggs and the mortality rate. A given queen can lay a certain maximum amount of eggs in optimal resource conditions and each landscape, with its particular distribution and density of resources, engenders a certain minimum degree of adult mortality. This minimum is due simply to the energy expenditures required to collect resources in that landscape. Together, the existence of a maximum birth rate and minimum death rate serve to limit population size, even if resources (pollen and nectar) are not limiting.

#### 3.1.2 Resource shortage

Secondly, we investigate the effect of resource shortage on colony growth and survival. We simulated the case in which the landscape is initialised with abundant pollen and nectar resources, and then one or both of these are abruptly and significantly reduced part way through the simulation. Fig. 3 shows how population size evolves before and after the induced shortage. A sudden shortage in nectar affects population size more dramatically than a sudden shortage in pollen, causing an exponential decay of the population that starts almost immediately after the nectar shortage occurs. The exponential decay occurs because nectar shortage increases mortality in addition to halting egg-laying. The response to the nectar reduction is rapid, because of the rapid decay rate of nectar stores through the energy costs of somatic maintenance, heating, and foraging. In contrast, a pollen shortage only causes a linear decrease in the population that starts some time after the pollen reduction. The decrease in population size is slower in the case of pollen reduction for two reasons. First, low pollen stores only halt births, but do not cause an increase in adult mortality, and second, pollen stores decay less quickly than nectar stores (pollen doesn’t evaporate and can be easily stored in large quantities).

**Figure 3:**
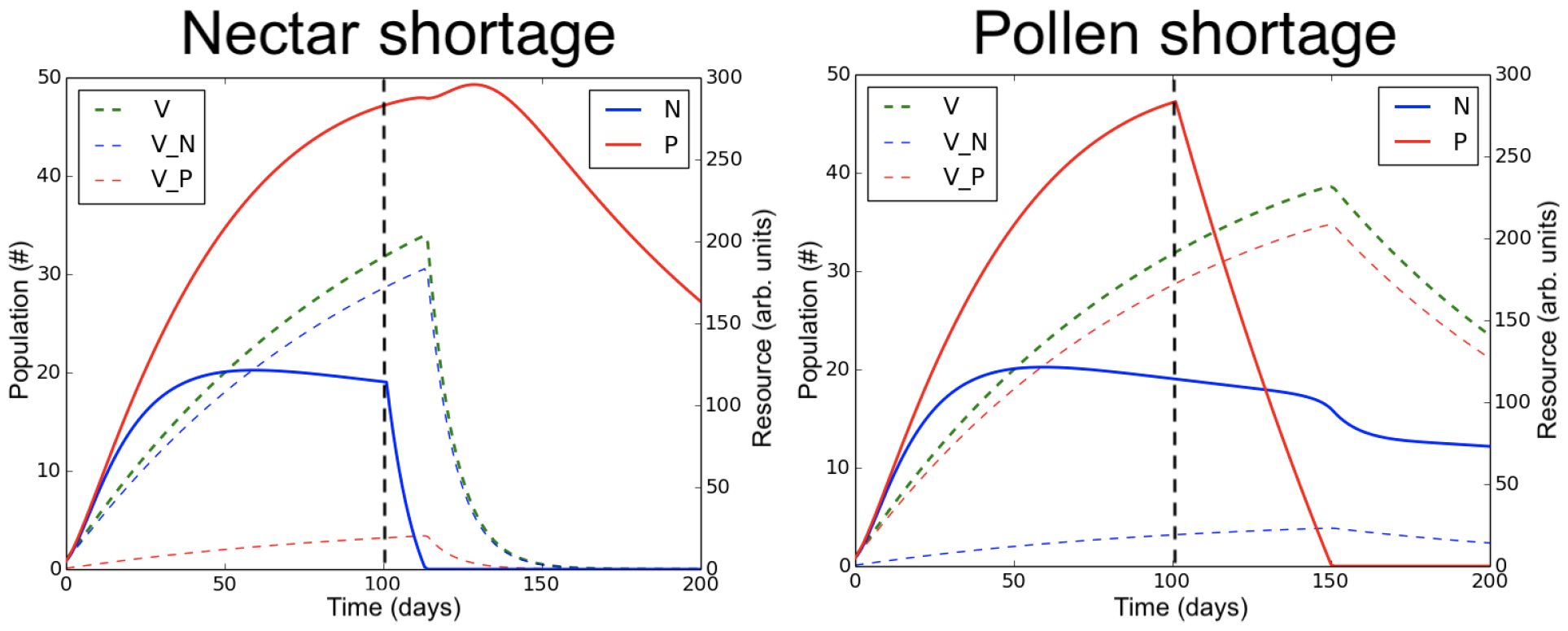
Structural energy of the colony (*V*) (stored in nectar foragers (*V*_*N*_) and pollen foragers (*V*_*P*_)), as well as nectar storage (*N*) and pollen storage (*P*) as functions of time, in a landscape with a uniform distribution of resource. On the left, we present the case in which the nectar resources in the landscape disappear at time 100 hours. On the right, the pollen resources in the landscape disappear at time 100 hours.

#### 3.1.3 Heating cost lead to extinction after mass crop bloom

In this section we investigate the effects of a mass crop bloom on colony survival. Fig. 4 shows three scenarios: A crop blooming in a landscape with (1) no, (2) medium, and (3) high nectar limitation. In all cases, we see that crop bloom represents the sudden addition of plentiful floral resources causing a rapid increase in colony size. When the crop bloom ends, the available resource level reverts to that of the original landscape. In all cases, colony size decreases exponentially, and, in the two nectar-limited landscapes, drops to extinction (or very nearly so). The exponential decrease is due to the increase in heating costs associated with a large nest, a mass crop bloom in a landscape otherwise very limited in nectar can lead to colony extinction.

**Figure 4:**
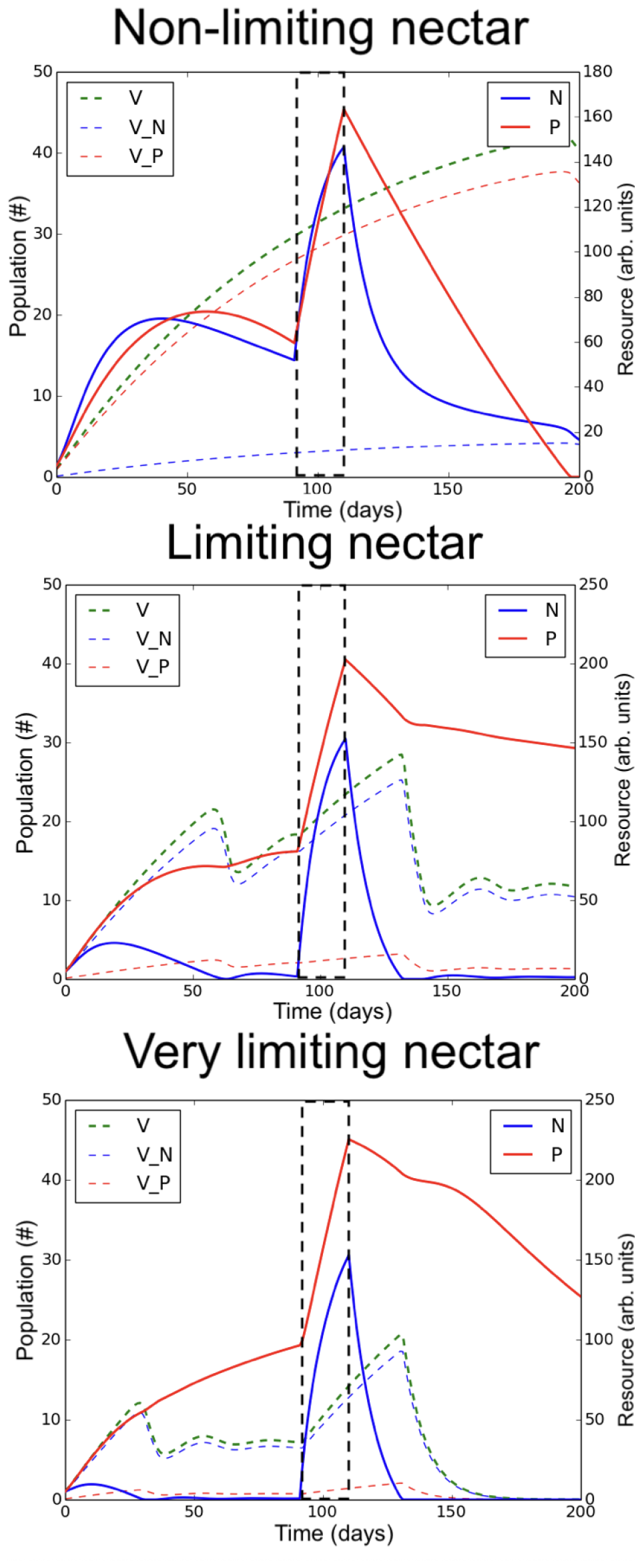
Structural energy of the colony (*V*) (stored in nectar foragers (*V*_*N*_) and pollen foragers (*V*_*P*_)), as well as nectar storage (*N*) and pollen storage (*P*) as functions of time, in a landscape with uniform distribution of resource. In the top plot, we represent the case in which the landscape has plentiful nectar resources (low agricultural intensity). In the bottom plot, we represent the case in which the landscape has very limited nectar resources (high agricultural intensity). In the centre, we represent an intermediate case where the nectar resources on the landscape are low but not severely so.

#### 3.1.4 Travelling costs

In order to understand the separate effects of distance to nectar resource and distance to pollen resource, we consider a simplified landscape containing a single nectar patch, a single pollen patch, and the bumble bee nest. The results are shown in Fig. 5. We see that increasing the distance to the nectar patch decreases colony population size more than increasing the distance to the pollen patch. We also observe oscillations in the population dynamics when the nectar patch is close to the nest but the pollen patch is some distance away.

**Figure 5:**
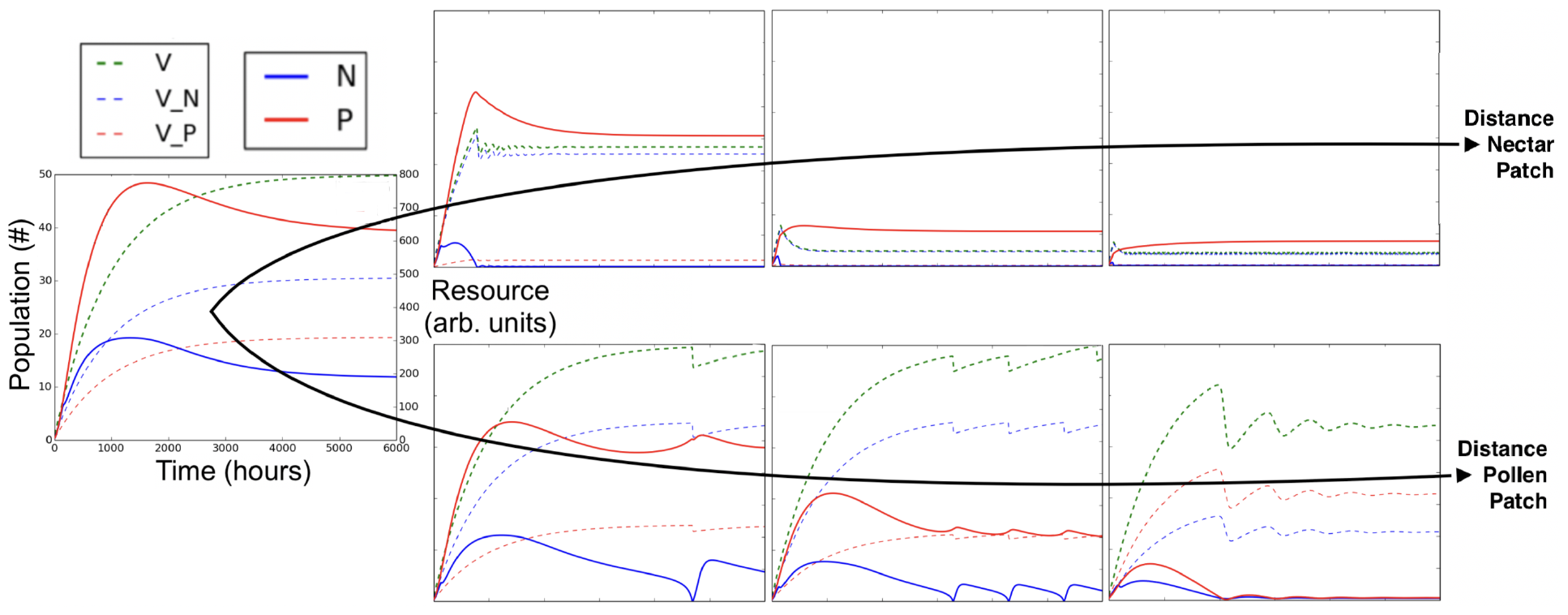
Structural energy of the colony (*V*) (stored in nectar foragers (*V*_*N*_) and pollen foragers (*V*_*P*_)), as well as nectar storage (*N*) and pollen storage (*P*) as functions of time, in a landscape containing one nectar patch and one pollen patch, each placed some distance from the nest. In the top row, the pollen patch is kept close to the nest site while the nectar patch is moved further away. In the bottom row, it is the nectar patch that remains close to the nest site while the pollen patch is moved further away.

#### 3.1.5 Benefits of planting wildflowers in nutritionally deficient crops

In Capera-Aragones, Foxall, et al. (2022), the authors present simulations results for the spatial distribution of foragers and the pollination services they provide to nutritionally deficient crops. Two key result are presented:

1. In crops that are nutritionally deficient, i.e., crop flowers with which bees can not fulfill their nutritional needs, adding nutritionally rich wildflowers can significantly increase the crop pollination services.
2. The crop fields that benefit most from wildflower enhancements are those that are found in landscapes characteristic of intense agriculture, i.e., landscapes where the background level of wildflowers is low.

In this section, we aim to simulate the same scenarios as in Capera-Aragones, Foxall, et al. (2022), to see if we can recover their results using our very different mathematical approach. We therefore perform simulations in which we keep the population of bees constant through time (as the previous model does not consider population dynamics), and we assume pollination services to be proportional to the amount of resource collected by foragers, be that pollen or nectar. In addition, similarly to Capera-Aragones, Foxall, et al. (2022), we define the relative nutritional composition as a percentage given by:

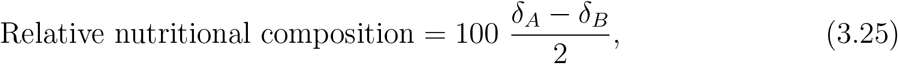

where

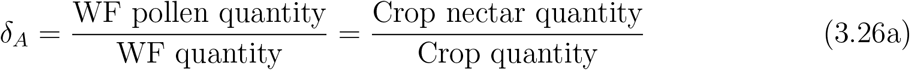

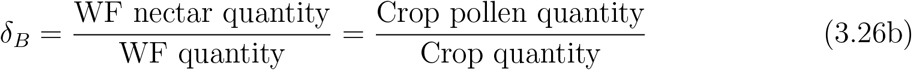

and *δ*_*A*_ and *δ*_*B*_ range between 0 and 2 and satisfy *δ*_*A*_ + *δ*_*B*_ = 2. With definition, ‘relative nutritional composition’ of 100% means that the wildflower resources are completely different in pollen and nectar composition from the crop resources, while a wildflower differentiation of 0% means that the compositions are exactly the same.

Fig. 6 shows the relative change in crop pollination services as the relative nutritional composition of pollen increases, for three different quantities of wildflowers planted. In Fig. 6 we see that the increase in crop pollination services is higher for high relative nutritional composition of pollen. In addition, we see that large quantities of wildflowers are most beneficial, but only when the relative nutritional composition is high.

**Figure 6:**
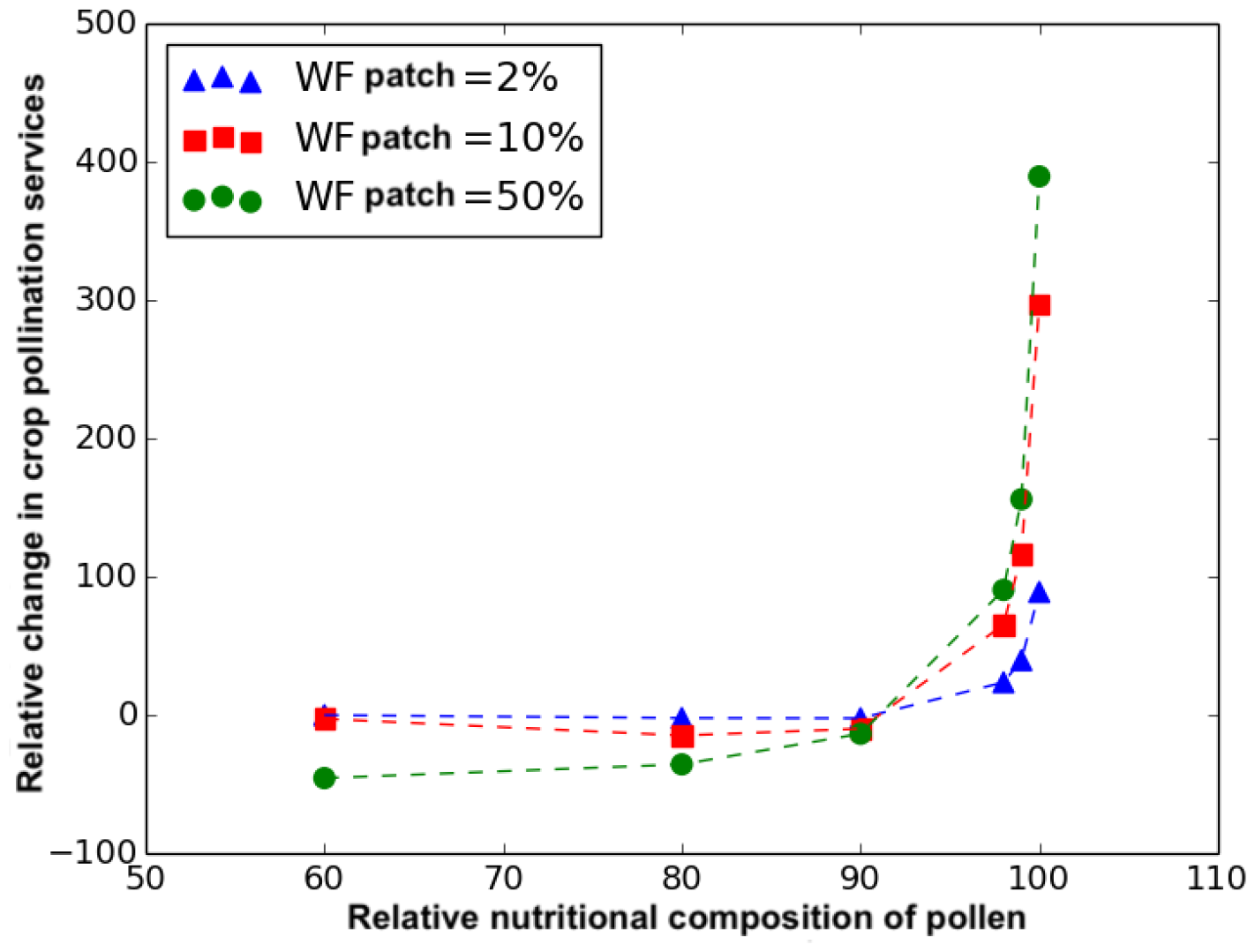
Percentage of increase in crop pollination services as a function of the relative nutritional composition of wildflowers and crop flowers, for three different quantities of wildflowers added (one in which the wildflower patch represents 2% of crop’s size, one of 10%, and one of 50%). The relative nutritional composition is a measure of the relative nutritive value of crop flower to wildflower pollen and is given by equation Eq. 3.25. The distance from the wildflower patch and the crop to the nest site are equal.

Fig. 7 shows how the benefits of planting wildflowers on crop pollination services increase as the agricultural intensity in the landscape increases. Agricultural intensity is inversely proportional to the background level of wildflowers in the landscape. The results for increasing wildflower patch size are nested, meaning that larger wildflower patch sizes are more beneficial across all levels of agricultural intensity, though particularly when background wildflowers are few.

**Figure 7:**
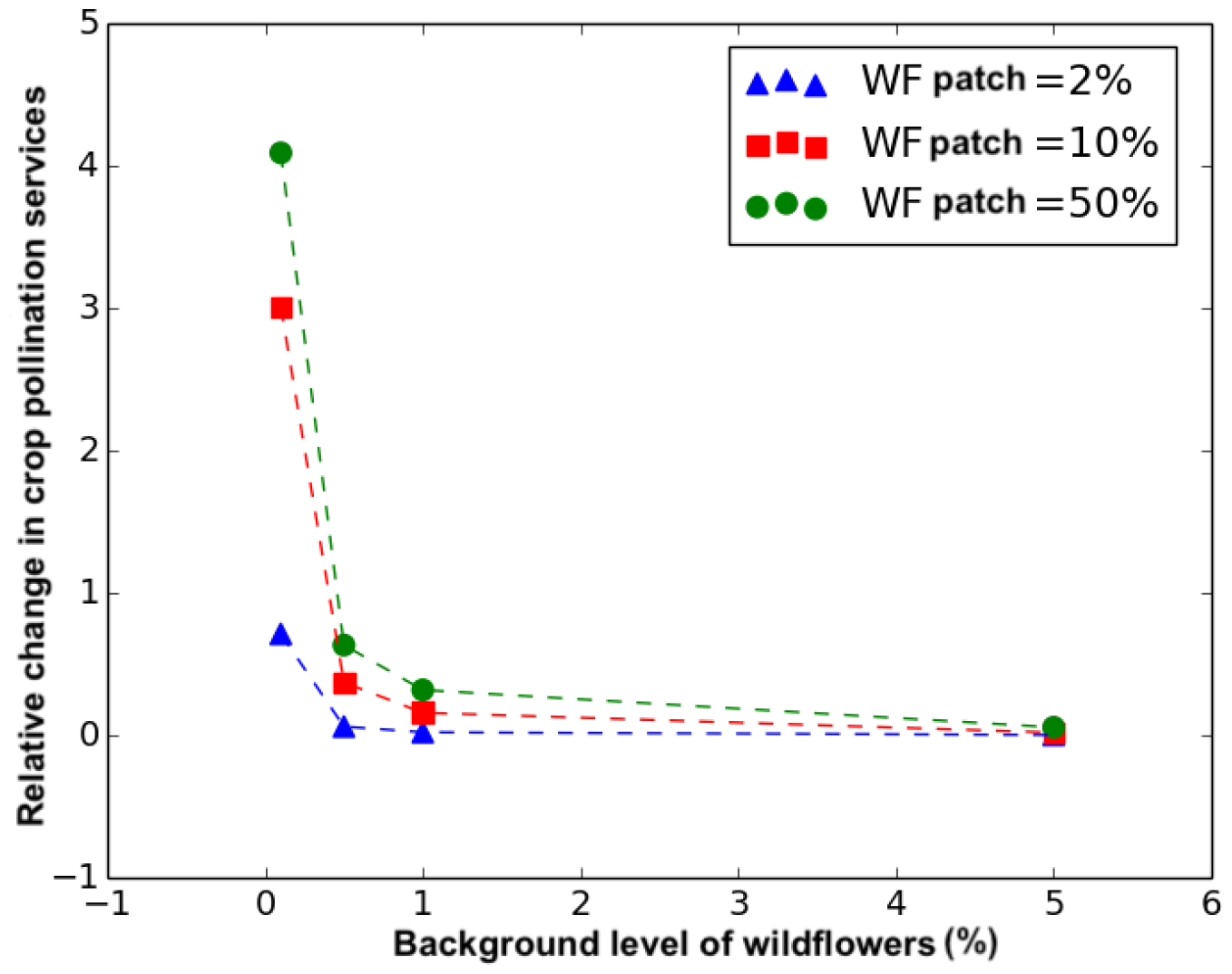
Percentage of increase in crop pollination services as a function of the background level of wildflowers in the landscape, which is inversely proportional to agricultural intensity. Results are shown for three different sizes of wildflower patches. The relative nutritional composition of wildflowers and crop flowers is set to 100% for this simulation.

### 3.2 Multiple seasons and multiple colonies

#### 3.2.1 Planting wildflowers can increase colony size

Blaauw and Isaacs (2014) published an empirical study showing that the addition of wildflowers adjacent to a blueberry crop can increase the population of wild pollinators, and the pollination services to the crop. Here we aim to reproduce these results.

In Blaauw and Isaacs (2014) they planted a wildflower patch 30% the size of the crop area, and the wildflowers bloom from May to October. Although the size and blooming time of the wildflower patch is specified, the density of flowers as a function of time is not. Therefore, to reproduce the results in Blaauw and Isaacs (2014) using our model, an assumption on the time evolution of the density of flowers is required. In this work we assume that the wildflowers add 30% of crop resource density all over the season. We also need to make assumptions about the location of the wildflower patch relative to the colony, the relative nutritional composition, and the intensity of the agriculture in the local landscape (i.e., the background level of wildflowers). Here, we assume that the wildflowers are located in the best place possible (previous studies suggested that locating the wildflowers on the far edge of the crop with respect to the nest site is consistently a good location (Capera-Aragones, Foxall, et al., 2021, 2022)), and we will study different values for the relative nutritional composition and the agricultural intensity.

In Fig. 8 we show the increase in crop pollination services as a function of the year since the addition of the wildflower patch. The plot shows two different relative nutritional compositions and three levels of wildflower background densities. We can see that, over the years, adding wildflowers becomes increasingly beneficial, especially in agriculturally intense crops (with a low wildflower background) and when the relative nutritional composition is high. We can also see in Fig. 8 that the increase of the population tends to saturate over the years for a low relative nutritional composition, as it is also observed in Blaauw and Isaacs (2014). For a high relative nutritional composition, the saturation also occurs but takes longer (tested using simulations but not shown in the plot).

**Figure 8:**
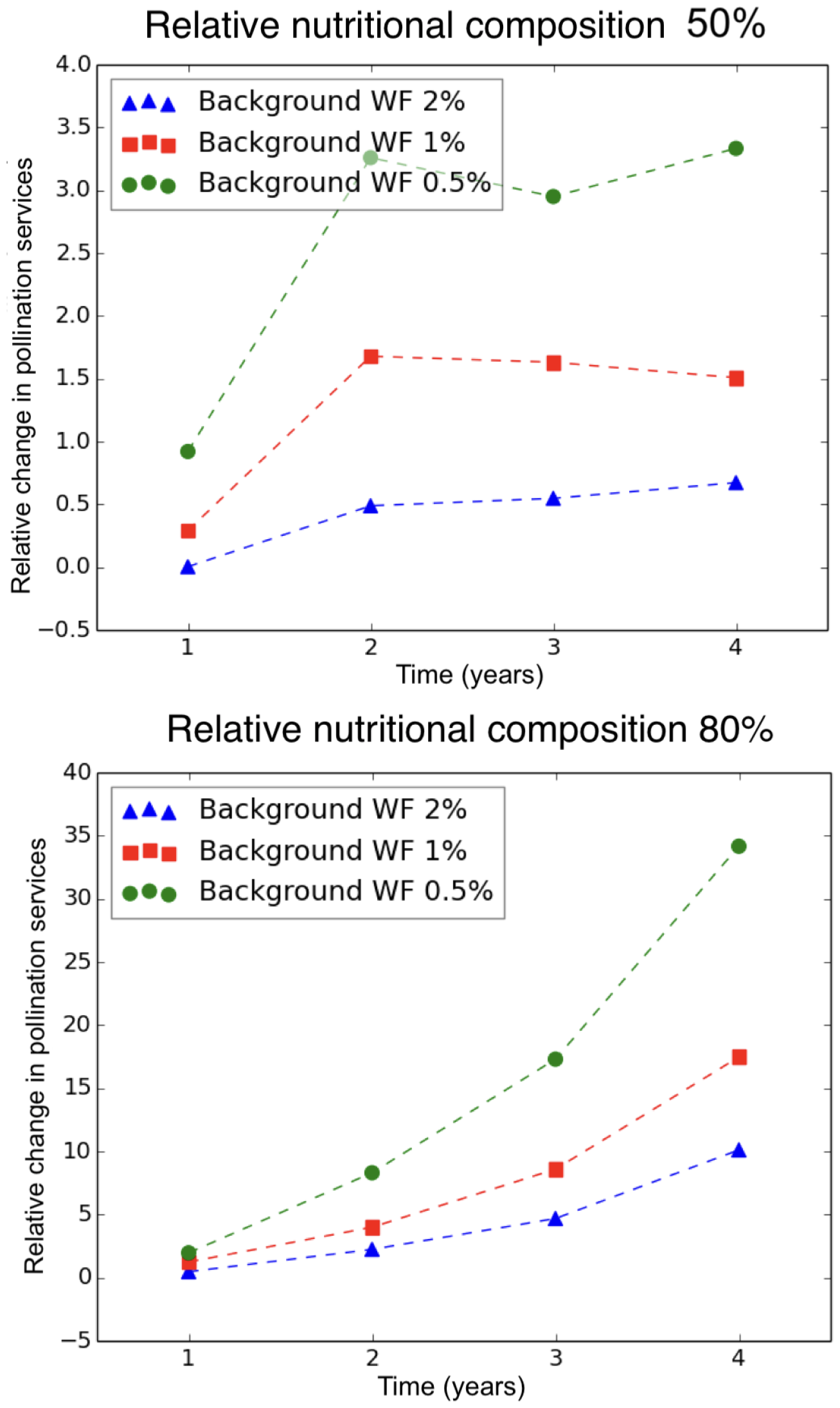
Relative increase in the crop pollination services as a function of years since establishment of the wildflower patch enhancements for two different relative nutritional composition (50% and 80%) and three wildflower background densities (or agricultural intensities). The two subplots have vertical axes with different scales. Wildflower patch was set to produce 30% of crop’s resource density. The bloom time for the crop was set from April 15th to 30th, the bloom time for the wildflower patch and for the wildflower low density background was set from April 1st to October 1st.

#### 3.2.2 Finding the best wildflower patch enhancement for crop pollination

In this section, we want to find the characteristics of the wildflower patch that would most benefit crop pollination services. To do so, we want to answer the five following questions:

1. Where to plant the wildflowers? (determine the location).
2. How many wildflowers? (determine the quantity or density).
3. Which relative nutritional composition should the wildflowers have? (determine the type of wildflowers depending on their pollen/nectar composition).
4. When should the wildflowers bloom? (determine the bloom time).
5. Does the level of agricultural intensity change the results? (determine if changing the wildflower background causes qualitative changes on the predictions).

In Capera-Aragones, Foxall, et al. (2021, 2022) and MacQueen et al. (2022), using mechanistic spatial models, they show that the best location for the wildflower patch is consistently the side of the crop farthest from the nest. For that reason and for simplicity of our analysis, in all the simulation of this section, we run simulations consistent with this scenario. That is, we set the distance between the nest and wildflower patch equal to the distance between the nest and furthest point in the crop field.

In addition, in this section we will limit our analysis to one level of agricultural intensity. Our results in Fig. 7 and Capera-Aragones, Foxall, et al. (2022) indicate that pollination services decrease monotonically with agricultural intensity, and so the qualitative patterns evident in the results below apply across agricultural intensity levels.

Fig. 9 shows how changing the three remaining characteristics of the wildflower patch (bloom time, wildflower type, and quantity) change crop pollination services. First, with respect to bloom time, the results clearly show that planting wildflowers that bloom both before and after crop bloom is by far the best strategy. This result can be explained as follows: wildflower resources that are available before crop bloom ensure that more bees are available to pollinate once crop bloom begins, and wildflower resources that are available after crop bloom ensure that the colony continues to grow and is able to produce queens (which will hibernate over the winter and initiate new colonies the following spring). The magnitude of the effect, however, is remarkable; wildflower patches that bloom ‘before and after’ crop bloom are significantly better than, the other cases investigated, that is, wildflower patches that bloom only ‘before’, ‘during’, or ‘after’ crop bloom.

**Figure 9:**
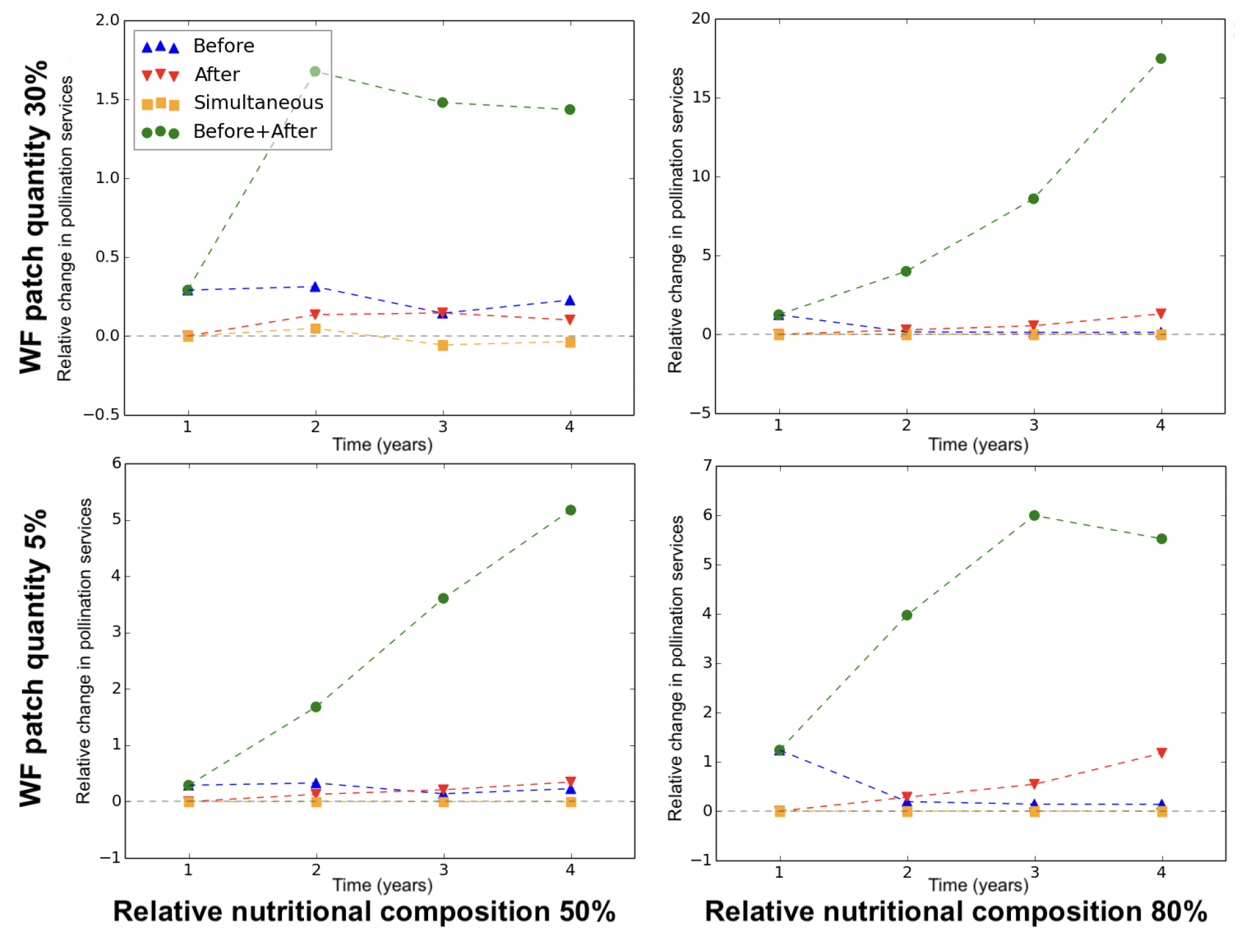
Increase in crop pollination services relative to the case when no wildflowers are planted, as a function of the years since patch introduction. Results are plotted for four different bloom times of the wildflowers relative to crop bloom, wildflowers blooming before (blue), after (red), simultaneously with (orange), and both before and after but not simultaneously with (green) the crop. The results are plotted for two different wildflower patch quantities (30% (top row) and 5% (bottom row)), and two different values of wildflower relative nutritional composition (50% (left column) and 80% (right column)). The agricultural intensity was set at a background level of wildflowers of 0.5%

Second, still looking at Fig. 9, we see that a higher relative nutritional composition is always beneficial for crop pollination services. Note that high differentiation is only possible for nutritionally deficient crops for which the wildflowers can provide complementary nutritional resources.

Third, we see that the optimal quantity of wildflowers depends on the relative nutritional composition (note the differences in the scaling on the vertical axes). In particular, when relative nutritional composition is high, a high density (quantity) of wildflower resource in the wildflower patch is best (compare left subplots), whereas, when relative nutritional composition is low, a low density (quantity) of wildflower resource is better (compare right subplots).

## 4 Discussion

In this work, we develop a relatively simple model framework to predict population dynamics, spatial distribution dynamics, and the interaction between the two. We then use the model to predict the coupled spatial and population dynamics of multiple wild bee colonies across several years.

### 4.1 A new modelling approach

To model the spatial distribution, we use the Maximum Entropy Principle (MaxEnt) as proposed by Capera-Aragones, Tyson, et al. (2023). This approach is computationally less intensive than current methods used to predict spatial distributions in that (1) the number of individuals considered does not affect simulation performance as in individual-based models, and (2) the spatial scale considered does not affect simulation performance as in partial and integro-differential equation models.

To model population dynamics, we develop a novel approach based on the application of a Dynamic Energy Budget (DEB) to an entire colony of bees, which is a departure from the usual approach of modeling individuals (Nisbet et al., 2012). The coupling of the colony-level DEB model with the spatial model given by MaxEnt is also novel, and allows us to efficiently explore the interplay between population dynamics and spatial distributions of foraging organisms.

In the first part of the paper, we use bumble bee colonies as our sample organism to demonstrate the effectiveness of our new modelling approach. We show that the model predicts reasonable wild bee colony behaviours in a diverse array of scenarios, e.g., changes to the colony growth rate when the landscape (without crop) is pollen- or nectar-limited, shrinking of the colony after a sudden shortage of resource, myriad effects ensuing from the temporary mass bloom of a crop, or the effect of travel costs on colony survival. The entire model, including the spatial component, consists only of ordinary differential equations, and so obtaining model solutions is straightforward and efficient.

### 4.2 Pollination services from wildflower patches: Results from our new modelling approach

In the second part of the paper, we ask how the addition of wildflower patches near crops affect crop pollination services over multiple years. Empirical studies are contradictory (Haaland et al., 2011; Lander et al., 2011; Nicholson et al., 2019; Sidhu and Joshi, 2016), with most measuring wild bee abundance rather than pollination services. Consequently, aspects of this same question have been the focus of a number of theoretical studies. When considering the period of a single crop bloom and using spatially explicit models, these studies show that wildflower patch location matters at the scale of individual foraging bees (MacQueen et al., 2022) and at the scale of the population (Capera-Aragones, Foxall, et al., 2021) when memory is included as a factor affecting foraging behaviour. Memory in these studies consists of repetitive returns to a chosen foraging site (MacQueen et al., 2022), or a process of learning and forgetting where the learning process uses the population density of foraging bees as a proxy for the density of foraging sites in memory (Capera-Aragones, Foxall, et al., 2021). The location effect is amplified when the crop floral resources are deficient in some way, and the wildflower patch resources compensate for this deficiency (Capera-Aragones, Foxall, et al., 2022). If the season between the production of the first cohort of workers and the start of crop bloom is also included, a recent spatially implicit model shows that wildflower patches blooming before but not simultaneously with the crop provide the most benefit to crop pollination services (Carturan et al., 2023).

Häussler et al. (2017) present a two-season and multi-annual model of wild bee population dynamics, coupled with a simple model of bee patch visitation and pollination services, and in landscapes typical of southern Sweden where habitat patches tend to be relatively small and divided by hedgerows. They find that late-blooming flower strips outperform grassy field margins, even though the former compete with crop flowers for wild bee visits. The results, however, depend on the season (early or late summer) when the results are measured. Memory-guided behaviour is not included in this model, but resource preference is.

In the present paper, for the first time, we combine the benefits through spatial dynamics, population dynamics, and the nutritional balance between crop flowers and wildflowers to determine the wildflower patch type that most increases crop pollination services. Our results, in agreement with (Capera-Aragones, Foxall, et al., 2021, 2022), show that when it is possible to plant wildflowers with floral resources that complement those offered by the crop flowers, the wildflowers should be planted in large quantities (or at high density). In contrast, when the relative nutritional composition of crop flowers and wildflowers is similar, then it is still beneficial to plant wildflowers, but in smaller quantities (or at lower density). Our model also shows, that agricultural intensification increases the need for wildflower patches, confirming earlier observations (Deguines et al., 2014). With respect to bloom time, we find that the best wildflower patches bloom both before and after crop bloom, but not during crop bloom.

The results we present here are chiefly qualitative, showing that wildflower patches with the appropriate composition, bloom time, and placement can potentially have a large beneficial effect on crop pollination services. We anticipate that with future field measurements of key model parameters, the accuracy of the quantitative predictions could be improved. In particular, parameter measurements for specific locations and bee species would be helpful for local growers. A benefit of the Dynamic Energy Budget approach is that the parameters are directly linked to measurable colony properties. The parameters of the MaxEnt model are more theoretical, but conceptually straightforward and easily varied in simulation studies.

### 4.3 Summary

Our work provides a new framework coupling population dynamics, using a novel whole colony Dynamic Energy Budget approach, to a spatial distribution model based on the maximum entropy principle. This coupling gives modelers a valuable new tool for modelling spatio-temporal population dynamics. We demonstrate the power of our approach, using the wild be colony as our example oganism to study the effect of added wildflower patches on crop pollination services. Our results are consistent with earlier work, and provide new insights into the relationship between wildflower patch effectiveness and (1) agricultural intensity at the landscape scale and (2) nutritional differentiation between the floral resources of crop flowers and wildflowers.

## Acknowledgements

RCT acknowledge NSERC STPGP 506922-17 and NSERC DG RGPIN-2016-05277 grant. Also thanks BRAES and the BC Blueberry Council. EF acknowledge NSERC Discovery Grant. PC acknowledge Ralph Cartar for insightful discussions about bumble bee behaviour.

